# Characterization of the nuclear and cytosolic transcriptomes in human brain tissue reveals new insights into the subcellular distribution of RNA transcripts

**DOI:** 10.1101/2020.04.08.031419

**Authors:** Ammar Zaghlool, Adnan Niazi, Åsa K. Björklund, Jakub Orzechowski Westholm, Adam Ameur, Lars Feuk

**Affiliations:** Department of Immunology, Genetics and Pathology, Uppsala University; Science for Life Laboratory in Uppsala, Uppsala University; Department of Cell and Molecular Biology, National Bioinformatics Infrastructure Sweden, Science for Life Laboratory, Uppsala University, Husargatan 3, SE-752 37 Uppsala, Sweden; Department of Biochemistry and Biophysics, National Bioinformatics Infrastructure Sweden, Science for Life Laboratory, Stockholm University, Box 1031, SE-17121 Solna, Sweden

**Keywords:** RNA sequencing, transcriptome, nuclear RNA, cytosolic RNA, lncRNA

## Abstract

Transcriptome analysis has mainly relied on analyzing RNA sequencing data from whole cells, overlooking the impact of subcellular RNA localization and its influence on our understanding of gene function, and interpretation of gene expression signatures in cells. Here, we performed a comprehensive analysis of cytosolic and nuclear transcriptomes in human fetal and adult brain samples. We show significant differences in RNA expression for protein-coding and lncRNA genes between cytosol and nucleus. Transcripts displaying differential subcellular localization belong to particular functional categories and display tissue-specific localization patterns. We also show that transcripts encoding the nuclear-encoded mitochondrial proteins are significantly enriched in the cytosol compared to the rest of protein-coding genes. Further investigation of the use of the cytosolic or the nuclear transcriptome for differential gene expression analysis indicates important differences in results depending on the cellular compartment. These differences were manifested at the level of transcript types and the number of differentially expressed genes. Our data provide a resource of RNA subcellular localization in the human brain and highlight differences in using the cytosolic or the nuclear transcriptomes for differential expression analysis.

## Introduction

The mammalian transcriptome harbors a myriad of coding and non-coding RNA transcripts, with biogenesis, abundance and subcellular localization tightly regulated to match the physiology and function of the cell. Genomewide technologies such as RNA sequencing have been instrumental in characterizing the transcriptome architecture. However, transcriptome analysis using RNA sequencing has primarily been performed on total or polyA+ RNA from whole cells, overlooking the spatial dimension of gene expression at the subcellular level. Nevertheless, an increasing number of reports are pointing to the importance of investigating the subcellular repertoire of RNA molecules and understanding the mechanisms that govern their distribution inside the cell ^1–4^. Subcellular RNA localization is widespread and conserved from bacteria to mammals, and it is becoming evident that this process plays crucial roles in regulating gene expression ^5,6^. For mRNA, subcellular localization provides a means to spatially control protein production and target proteins to their site of function ^7–11^. Therefore, the differential distribution of mature mRNA between the nucleus and the cytoplasm may have a direct impact on protein expression levels. In fact, it has been recently shown that nuclear retention of mature mRNA is a mechanism that involves a wide range of protein-coding transcripts. This nuclear retention is believed to ultimately fine-tune mRNA translation in the cytoplasm ^12^. Similarly, identifying the subcellular localization of non-coding RNA such as long non-coding RNAs (lncRNAs) can provide substantial insights into their biology and function. Based on the link between localization, function, and regulation it is important to map the subcellular localization of coding and non-coding RNA in different cells and tissues to obtain a more comprehensive understanding of RNA’s biological functions.

Early attempts to determine the cellular localization of RNA molecules were performed on one transcript at a time, mostly using RNA fluorescent *in situ* hybridization (RNA FISH) ^13,14^. However, in the last few years, the field has benefited from the use of various high-throughput technologies, such as expression microarrays and RNA-seq in combination with cellular fractionation ^11,15–17^. The general aim of these studies was to characterize the contribution of each of the nuclear and cytoplasmic transcriptomes to the global gene expression profiles. Several studies have demonstrated that although the nucleus and cytoplasm contain overlapping populations of RNA transcripts, there are significant differences between the two fractions, and many coding and non-coding transcripts are unique to one fraction ^17^. The differentially localized transcripts exhibit particular global characteristics. For example, transcripts localized to the cytoplasm tend to have shorter coding sequences and shorter UTRs compared to the transcripts localized in the nuclear fraction^15,18^.

The subcellular expression patterns of lncRNA have also been the focus of several studies during the last years. Although a large number of lncRNAs have been identified, the biological relevance for the majority of lncRNAs is still elusive ^19,20^. Using microarrays, RNA-seq and imaging technologies, several reports have demonstrated that although lncRNA are mostly localized in the nucleus, cytoplasmic lncRNAs also exist ^19,21–25^. In fact, lncRNA subcellular localization was divided into three categories: 1. Exclusively expressed in the nucleus, 2. Mainly enriched in the nucleus and, 3. Mainly enriched in the cytoplasm ^26^.

Due to the importance of expanding the catalog of RNA subcellular localization, Zhang et al., established a database (RNALocate), from previously published data, that documents the subcellular localization of different RNA types in several species ^3^. However, most of the current knowledge about RNA localization is based on cell lines and in most cases restricted to targeted experiments rather than analysis of the complete RNA co-expression profiles. Global profiling of the differences between nuclear and cytosolic fractions is crucial for accurately interpreting data from single-cell transcriptome studies. Single-cell analysis from frozen tissue mainly involves sequencing RNA from sorted nuclei ^27^. Therefore, understanding the main differences between the nuclear and cytosolic transcriptome in human tissues is important for meaningful interpretation of results from single-cell RNA sequencing experiments.

In this study, we utilized an efficient cytoplasmic and nuclear RNA extraction protocol ^28^. to purify populations of subcellular RNA fractions from human fetal and adult frontal cortex, as well as fetal cerebellum. We investigate the differences in relative transcript abundance between cytosol and nucleus and highlight genes and gene categories that are over-represented in either compartment. Our results also provide important insight into the subcellular localization of nuclear-encoded-mitochondrial proteins (NEMPs) and noncoding RNA in brain tissue samples.

## Results

To characterize the abundance of transcripts in the nuclear and cytosolic RNA fractions from brain tissue we sequenced total RNA extracted from six adult frontal cortex samples, three fetal frontal cortex samples, and cerebellum from the same three fetal samples. The nuclear and cytosolic RNA was also sequenced from the human neuroblastoma cell line (SHSY-5Y) for reference. Total RNA sequencing was carried out using IonProton whole transcriptome with an average of 80.5 million mapped reads per nuclear sample and 46.9 million reads per cytosolic sample. The cytosolic sequence reads mapped primarily to exons (84.1% exonic mapping) while the nuclear reads, as expected, had a large proportion aligned to intron and intergenic sequences (19.81% exonic mapping). The lower percentage of exonic reads in the nucleus compared to the cytosol was due to the presence of nascent transcripts. Details about the samples and sequencing of each sample are provided in Supplementary Table 1. Hierarchical clustering of all samples based on the expression of protein-coding genes and lncRNAs showed brain tissues clustering together with their corresponding subcellular fraction (a cytosolic cluster and a nuclear cluster), while both fractions from the SHSY-5Y cell line formed a separate cluster (Figure 1A). Similarly, principal component analysis (PCA) separated all the samples according to their subcellular fractions and tissue sources along the PC1 and PC2, respectively (Figure 1B).

**Figure 1:**
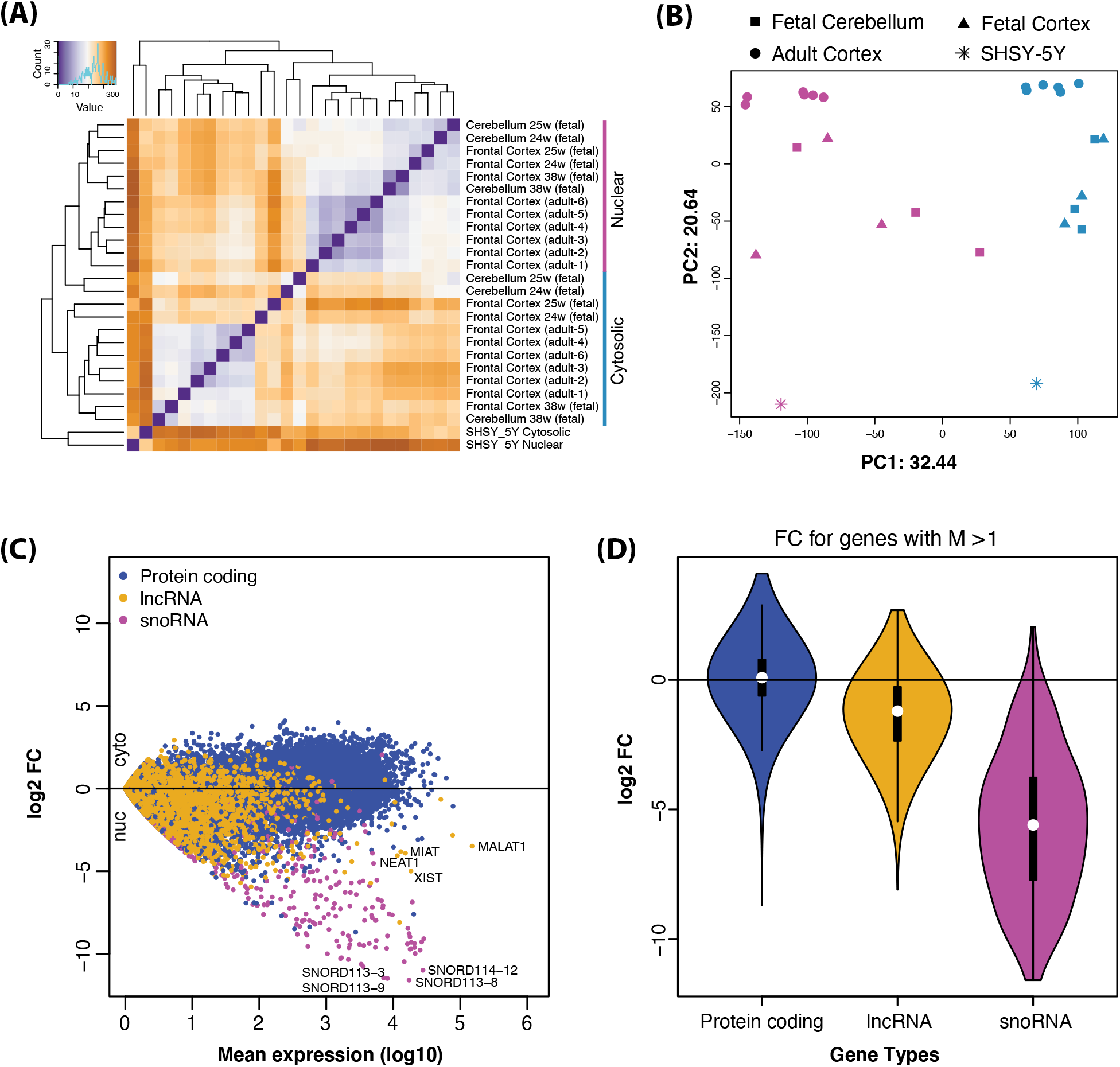
(A) Hierarchical clustering analysis of cerebellum (fetal), frontal cortex (fetal and adult) and SHSY-5 cytosolic and nuclear samples using DESeq2 rlog-normalized counts. The Euclidean distance values used for clustering are represented by a color code from brown (low correlation) to purple (maximum correlation). (B) Principal Component Analysis performed using the same rlog-normalized expression data. Contribution to variance for principal components, PC1 and PC2, are reported in the graph. (C) Expression means versus log_2_ fold changes (log_2_FC) showing global (all tissues) cytosolic and nuclear expression differences for protein-coding, lncRNA and snoRNA genes. Each point with log_2_FC > 0 corresponds to cytosolic localization whereas log_2_FC < 0 corresponds to nuclear localization of the genes. (D) Violin plot of gene biotypes in (A) displays log_2_FC (y-axis) distribution between cytosolic and nuclear fractions. White dots indicate mean, box edges represent the interquartile range, and the colored region and curve show the probability density function. Each group is composed of genes with log_10_ mean expression higher than one (M>1), and shown in similar colors as in (A).

To further investigate the distribution of transcripts across the cytosolic and nuclear fractions and identify genes that show differential abundance between nucleus and cytosol, we first investigated how different classes of transcripts are distributed. We found that protein-coding genes were equally distributed between the nuclear and the cytosolic fractions, whereas lncRNAs and small nucleolar RNAs (snoRNAs), as previously shown ^29^, were more abundant in the nuclear fraction (Figure 1C and D). We could confirm the nuclear localization of a number of highly expressed lncRNAs with known nuclear localization (including *XIST* and *MALAT1*). We observed similar distribution patterns when each brain tissue was analyzed separately (Supplementary Figure 1). However, the most skewed distribution was exhibited by snoRNAs, which are well-known to have their primary function in the nucleus. Examples of snoRNA with extreme nucleus bias include SNORD114-12 and SNORD113-8 (Figure 1C). Overall, the PCA and distribution of the different transcripts are in agreement with previous findings and indicate that our analysis readily identifies transcripts with a previously demonstrated cytosolic or nuclear localization^33^.

### Protein coding genes

To identify transcripts exhibiting differential distribution in the cytosolic and nuclear fractions, we first analyzed protein-coding transcripts across all tissues. In total, we identified 5,109 transcripts that were significantly more abundant in the cytosol and 5,397 transcripts significantly more abundant in the nucleus (*P_adj_*<0.05, Supplementary Table 2, Supplementary Figure 2A). A separate analysis grouped by tissue type and age showed more significant transcripts in both cytosol and nucleus in the adult samples, reflecting that the adult tissues represented a larger sample group with less inter-individual heterogeneity compared with the fetal tissues (Data not shown). To evaluate the potential contribution of nascent RNA to transcripts localized to the nucleus, we performed a similar analysis using polyA+ RNA-seq data from the cytosol and nucleus from the Encode cell lines. We found similar subcellular distribution patterns of transcript abundance seen in our brain cytosolic and nuclear total RNA-seq data (Supplementary Figure 2B-D). This indicates that nascent RNA does not fully explain the transcript localization observed in the nuclear fraction. Next, we performed the gene ontology (GO) enrichment analysis of genes that are significantly more abundant in cytosol and nucleus across all tissues. We found transcripts representing specific biological processes to be enriched (*P_adj_*<0.05). In cytosol, there was significant enrichment of transcripts involved in metabolic and catabolic processes, translation processes, macromolecular and protein complex organization and disassembly, and in the nervous system and neural projection development. Significant GO categories for cellular compartment showed that the biological processes associated with cytosolic transcripts can be primarily linked to the extracellular organelles, vesicles, and ribosomes. Whereas in the nucleus, there were fewer transcripts in a particular ontology and less number of significant GO categories in general. Significant categories include regulation of transcription, sensory processes, and regulation of various biosynthetic processes. See Supplementary Table 3 for lists of all enriched GO categories.

Previous studies demonstrated that transcripts localized to the cytosol or the nucleus possess particular genic features in terms of exon length ^17^. Analysis of transcript abundance differences between the cytosol and the nucleus and mature transcript length for the three sample groups indicates that long transcripts are more nuclear compared to short transcripts (Supplementary Figure 3A), highlighting a transcript length signature in subcellular mRNA distribution. However, no such pattern was observed when the transcript abundance differences between the cytosol and nucleus were analyzed against the UTR length of the genes (data not shown). We also found, when comparing the global expression levels between nucleus and cytosol, that genes with relatively higher expression tend to be shifted toward the cytosol (Supplementary Figure 3B). To examine whether the higher expression in the cytosol is caused by the transcript length bias detected between the cytosol and the nucleus, we analyzed the RPKM expression values and transcript lengths in the three sample groups. The results demonstrated no influence of transcript length on gene expression (Supplementary Figure 3C).

Although not evident in the GO analysis, when we manually investigated lists of differentially localized genes in the cellular fractions, we observed that many nuclear-encoded-mitochondrial proteins (NEMPs) were abundant in the cytosol. To further analyze the enrichment of NEMP transcripts, we extracted a full list of genes (1,158) encoding proteins with evidence for mitochondrial localization from Mitocharta2.0 ^34,35^. By investigating gene expression and fold change differences for NEMP transcripts in the cytosol and nucleus, we observed a clear shift for NEMP transcripts to the cytosolic compartment (Figure 2A) and that NEMPs were significantly enriched in the cytosol compared to all other protein-coding genes (p= 2.8×10e-76, Figure 2B). In our cellular fractionation protocol, we used a centrifugation power of 10,000 rpm, which results in the mitochondria being pelleted together with the nuclear fraction. Therefore, it is unlikely that the enrichment of NEMPs in the cytosol was due to NEMP RNAs being physically anchored to the mitochondria. We further validated this by showing that the transcripts produced by the mitochondria were clearly nuclear in their localization (p= 6.4×10e-08, Figure 2A and 2B) indicating that the mitochondria themselves indeed end up in the nuclear fraction. The results of preferential cytosolic localization of NEMPs and nuclear localization of mitochondrial transcripts were consistent across all the tissues (Supplementary Figure 4). To experimentally validate the cytosolic localization of NEMPs, we performed RNA Fluorescence *in situ* hybridization (RNA FISH) ^36^ on two highly expressed NEMP targets in SHSY-5Y cells. The results demonstrated that both of the NEMPs targets were preferentially localized in the cytosolic fraction (Figure 2C). To further test if the cytosolic enrichment of NEMPs was consistent across various tissue types, we extended our analysis to cytosolic and nuclear RNA-seq data from 11 cell lines from the ENCODE project ^22,37^. In agreement with our brain data, we found that NEMPs exhibited preferential cytosolic localization in all the cell lines (Supplementary Figure 5), indicating that the cytosolic localization of NEMPs is a general phenomenon that involves various human tissues.

**Figure 2:**
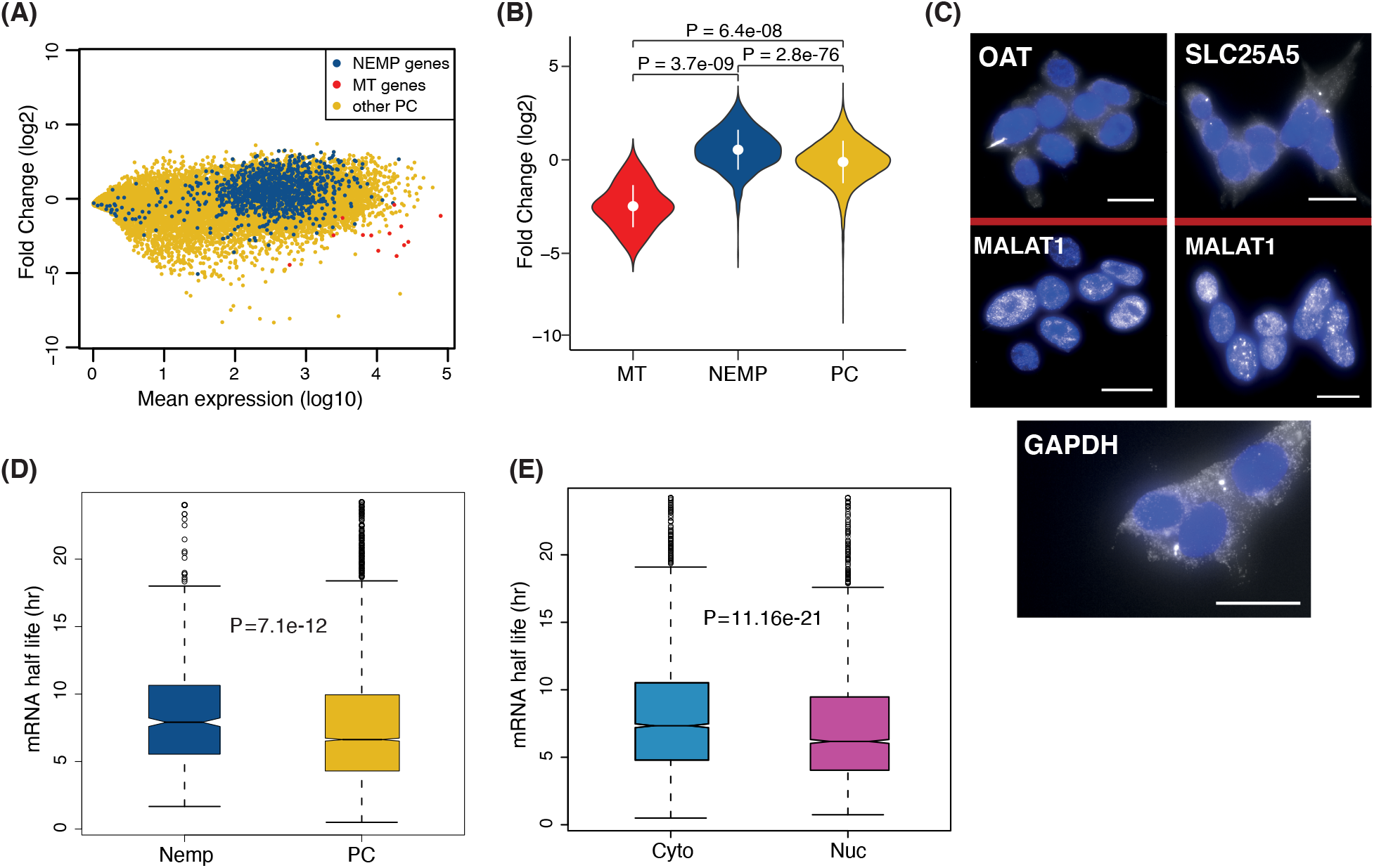
**(A)** MA plot showing localization distribution of NEMPs (dark blue), remaining protein-coding genes (yellow), and mitochondrial encoded genes (red). (B) Violin plot showing fold change distributions between the three gene categories (baseMean>=10) in cytosolic and nuclear fractions. P-values reported were obtained with the Mann-Whitney-Wilcoxon test. (C) Validation of the enrichment of NEMPs RNA transcripts OAT and SLC25A5 in the cytosol in SHSY-5Y cells using RNA FISH. MALAT1 (known nuclear transcript) and GAPDH used for comparison. Images represent maximum projections of 13, 18, 20 optical intervals spaced by 0.5 μm. Scale bar, 2 μm. (D) Box plot shows mRNA half-life rate comparison between groups of differentially localized NEMP (dark blue) and remaining protein-coding genes. (E) Comparison of mRNA half-life rate between cytosolic and nuclear DEGs.

One explanation for the increased abundance of a group of transcripts in the cytosol could be an increased mRNA half-life. To examine if the preferential cytosolic localization of NEMPs was due to long half-life, we used published mRNA half-life data ^38^ and inquired whether these transcripts have a longer half-life than the rest of protein-coding transcripts. We found that NEMPs displayed significantly longer half-life compared to other protein-coding transcripts (p= 7.1×10e-12, Figure 2D), which could explain the preferential enrichment of NEMPs in the cytosol. An analysis of the half-life of all proteincoding transcripts significantly associated with the cytosolic or nuclear compartments showed a significant difference in half-life, with a longer half-life for transcripts enriched in the cytosol (p= 1.6×10e-21, Figure 2E).

### Long non-coding RNA (LncRNAs)

We next investigated if lncRNAs also exhibit differential distribution in the nuclear and cytosolic fractions across all samples. Overall, we found a higher representation of lncRNAs in the nucleus (Figure 3A). Out of all expressed lncRNAs, we identified 1,114 lncRNAs in the nucleus compared to 118 lncRNA in the cytosol (*P_adj_* <0.05; Supplementary Table 4). The analysis in Figure 3A also shows the nuclear localization of previously reported nuclear lncRNAs including *MALAT1, MEG3*, and *XIST*. The nuclear and cytosolic enrichment of lncRNA in the individual tissues are shown in Supplementary Figure 6 (A-C). The adult frontal cortex samples harbored the highest number of lncRNAs that are significantly more abundant in the nucleus or the cytosol, followed by the fetal cortex and fetal cerebellum, respectively (Figure 3B). Although there was an overlap within the cytosolic and within nuclear lncRNAs between the tissues, each tissue type harbored a group of unique lncRNAs. We further investigated these tissue-specific cytosolic and nuclear lncRNAs, which demonstrated differential localization between the cytosol in a single tissue and found that the majority of these were either not expressed or expressed at low levels in the other tissues.

To further understand how subcellular localization of lncRNAs varies between different tissue types, we explored the subcellular distribution of lncRNAs in 10 cell lines from the ENCODE project. In this analysis, we only focused on lncRNAs that showed significant subcellular localization in either the cytosol or the nucleus in the brain samples. Although we found an overlap of lncRNAs between the subcellular compartments, there were many lncRNAs that exhibited differential subcellular localization between the brain samples and the cell lines (Figure 3C). A range from 35 to 69 lncRNA displayed differential localization between the brain and each of the different cell lines (Supplementary Table 5), such as *LINC00672*, which was cytosolic in the brain and nuclear in 7 out of 10 cell lines. The lncRNAs *RP11-452F19.3* and *RP11-449H11.1* were nuclear in the brain and cytosolic in 5 and 3 different cell lines, respectively. We also found that many of the lncRNAs that display differential localization between the brain and cell lines differ within the different cell types (Supplementary Table 5). These results further support that subcellular localization of lncRNAs is often tissue specific. The lncRNAs that displayed similar localization patterns between the brain and cell lines are listed in Supplementary Table 6.

**Figure 3:**
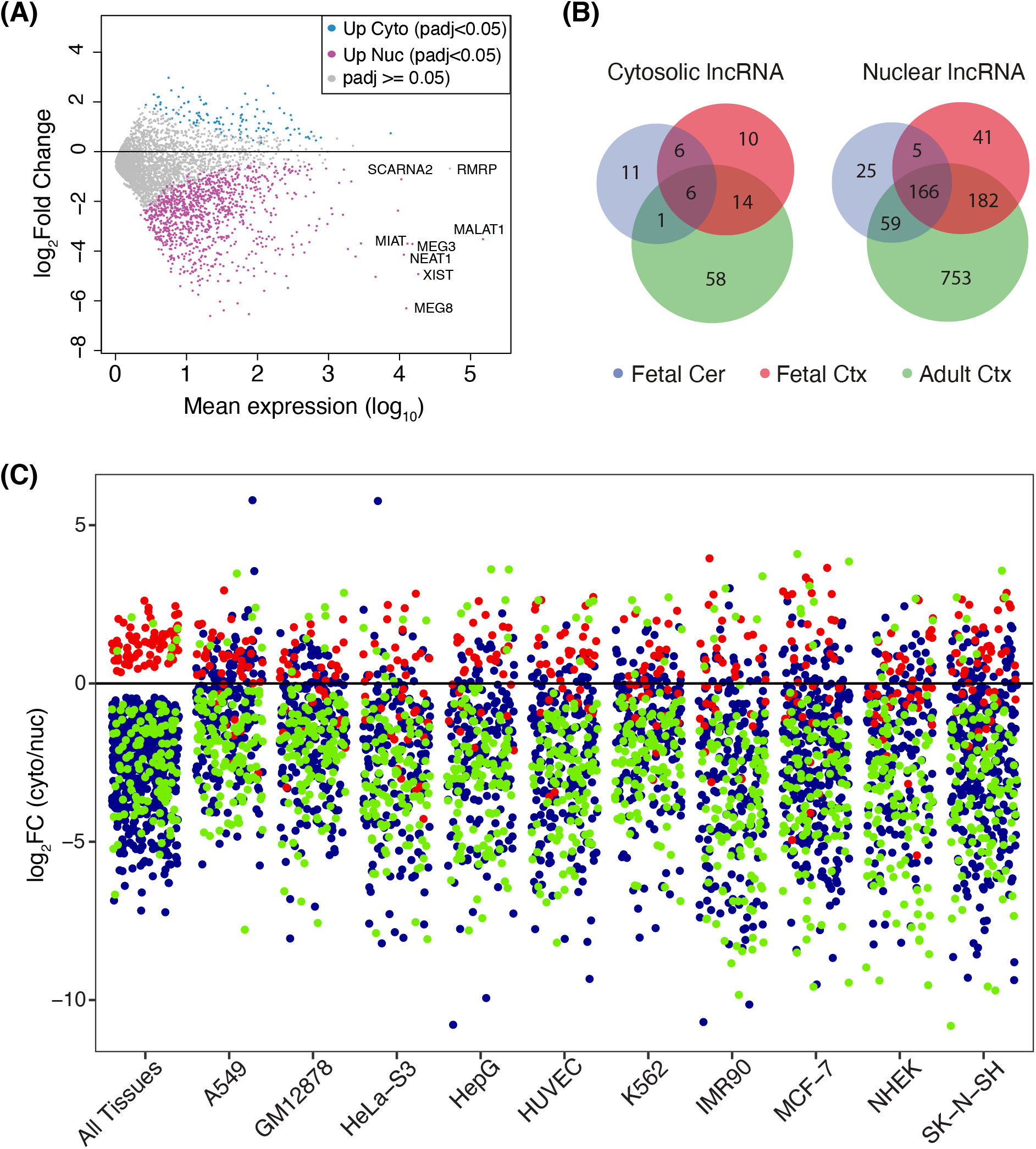
(A) MA plot showing global (all tissues) distribution of differentially localized lncRNA genes in the cytosol (blue) and nucleus (magenta). Nonsignificant lncRNA genes are highlighted as grey. (B) Venn diagram displaying overlapping lncRNA genes differentially localized in the fetal cerebellum (blue), fetal cortex (green) and adult cortex (red). (C) Strip chart showing localization of significantly cytosolic (red) and nuclear (blue) lncRNAs from our data (all tissues combined) and in ENCODE cell lines (baseMean >=10). Only differentially expressed lncRNAs in cell lines overlapping with alltissue DEGs are shown. Genes that remained consistently cytosolic or nuclear in the brain samples and cell lines are highlighted in green.

### The cytosolic and nuclear transcriptomes as a source for gene expression analysis

There is a longstanding interest in understanding the reliability of using the cytosolic or nuclear RNA as a proxy for whole-cell gene expression analysis. To obtain a more accurate view of the differences between the cytosolic and nuclear transcriptomes as a source for gene expression analysis, we compared the results of differential gene expression analysis between cytosolic and nuclear transcriptomes from the fetal frontal cortex and the adult frontal cortex (cytosolic vs cytosolic [cyto-cyto], and nuclear vs nuclear [nuc-nuc]). To control for the differences in the amount of sequencing data produced from the fetal and adult tissues (Supplementary Table 1), we first subsampled the read counts to obtain equal library sizes for the samples. Differentially expressed genes (DEGs) between the two samples for the cyto-cyto and nuc-nuc before and after subsampling showed that the subsampling had a marginal effect on the number of DEGs (Figure 4A). Subsampled data were used for the rest of the analysis in Figure 4. We then compared the number of up-regulated and down-regulated genes between the two tissues in the cyto-cyto and nuc-nuc data. There was a clear difference in the number of DEGs detected in cyto-cyto and nuc-nuc, with a higher number in the cyto-cyto analysis compared to the nuc-nuc (Figure 4B). Comparing the lists of DEGs between the two analyses revealed that although there was an overlap between the DEGs identified from the two analyses, a large fraction of the DEGs was unique to cyto-cyto analysis (Figure 4B). GO analysis for the unique list of DEGs obtained from the cyto-cyto comparison revealed enrichment of transcripts related to translational processes, various metabolic and biosynthetic processes and gene expression and RNA processing. Meanwhile, the unique DEGs from the nuc-nuc analysis were mainly enriched for transcripts that belong to mitochondrial translation, which as we mentioned was a result of the mitochondria being separated with the nuclear fraction in the extraction protocol (Supplementary Table 7-10). Dividing the lists of DEGs from Figure 4B into protein-coding and lncRNA revealed that even though the nuc-nuc analysis resulted in a smaller number of protein-coding genes, it harbored a larger number of differentially expressed lncRNAs as compared to the cyto-cyto analysis, which was not surprising given that most lncRNAs were localized to the nucleus (Figure 4C). Collectively, these results indicate that although the cytosol and the nuclear fractions contain largely overlapping lists of DEGs, each fraction also contains unique groups of DEGs. To our surprise, when we investigated the expression fold change of all DEGs obtained from cyto-cyto, and the nuc-nuc analysis, a group of 61 genes showed opposite differential expression patterns (Figure 4D). For example, *ETV2*, which encodes a transcription factor and is overexpressed and critically required during development, was up-regulated in the fetal sample when using cyto-cyto analysis and up-regulated in the adult sample when using the nuc-nuc analysis. Other genes that showed opposite differential expression patterns are highlighted in Figure 4D and listed in Supplementary Table 11 and Supplementary Table 12. These results imply that gene expression analysis using either of these fractions could lead to different conclusions on differential expression between samples. Therefore, we suggest that neither the cytosolic nor the nuclear transcriptomes alone represent an accurate proxy for a complete view of gene expression levels in a cell. In addition, using either the cytosol or the nucleus may lead to conflicting data.

**Figure 4:**
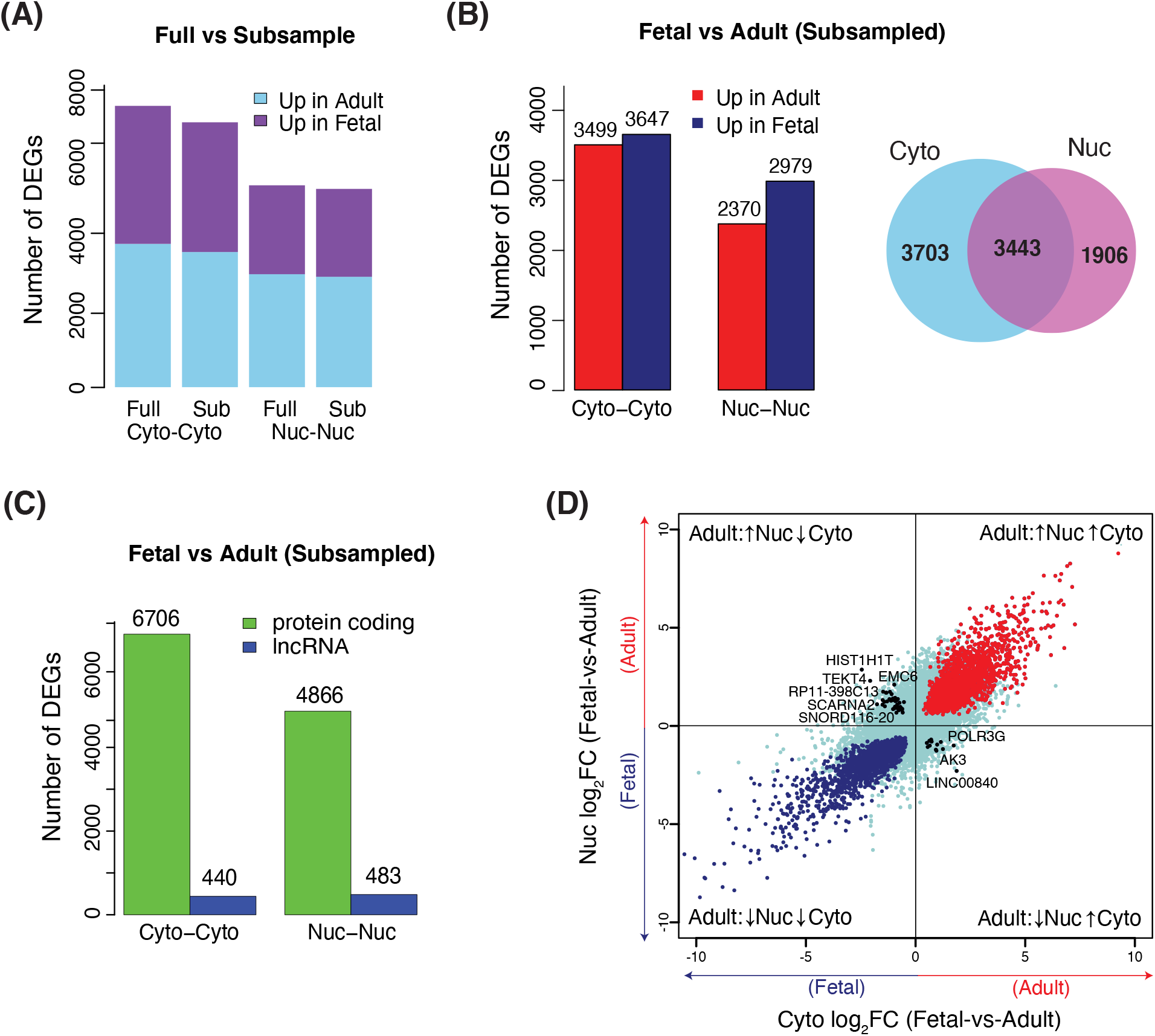
(A) Comparison of differential expression analysis between full and subsampled data. DEGs in Fetal-cyto vs. Adult-cyto and Fetal-nuc vs. Adult-nuc from the two datasets are shown. (B) Barplot showing DEGs in fetal and adult (cyto-cyto) and DEGs in fetal and adult (nuc-nuc). Overlap of total cytosolic and total nuclear DEGs is shown in the Venn diagram. (C) Barplot showing the distribution of differentially expressed protein-coding and lncRNA genes in cyto-cyto and nuc-nuc analysis. (D) Expression fold changes for transcripts in fetal and adult cortex cyto-cyto against nuc-nuc after subsampling. DEGs (red) overlap between cytosolic and nuclear fractions in adult, cytosolic and nuclear in fetal (blue). Genes significantly abundant in the fetal cortex using cytosolic samples but abundant in the adult cortex with nuclear samples, and those abundant in adult (cytosol) but not fetal (nuclear) are highlighted in black. Genes not reaching significance in either of the groups are shown in light blue.

A recent study by Lake *et al*. revealed a high concordance in the nuclear and whole-cell transcriptomes from mice when profiling cellular diversity using celltype-specific marker gene expression signatures ^39^. We compared our list of significant cytosolic and nuclear transcripts obtained from all brain samples with the whole-cell and nuclear data from Lake *et al*. and found that our analysis resulted in a larger number of cytosolic/nuclear-localized genes (5201 compared to 1518 cytosolic genes, and 5548 compared to 791). This is more likely due to the higher complexity and heterogeneity of samples used in our study. We also found that 65% of their cytosolic-accumulated and 45% nuclear-accumulated genes overlapped with our results, which validates the reliability of our analysis (Supplementary Figure 7). We investigated for overlaps between our list of genes that demonstrated opposite differential expression patterns between the cytosol and nucleus (Supplementary Table 11 and 12) with the list of cell-type-specific genes from Lake *et al*. We found three cell-type specific markers: *Hsd17b10* for Oligodendrocyte, *AK3* for Astrocyte, and *Chrnb1* for Endothelial cells which showed opposite differential expression patterns between the cytosol and the nucleus i.e. *Hsd17b10* and *Chrnb1* were upregulated in the fetal sample when using cyto-cyto analysis and upregulated in the adult sample when using the nuc-nuc analysis. Meanwhile, *AK3* was upregulated in the adult sample when using cyto-cyto analysis and upregulated in the fetal sample when using the nuc-nuc analysis.

## Discussion

To date, transcriptome analysis has mainly relied on analyzing RNA sequencing data from whole cells, overlooking the impact of subcellular RNA localization and its influence on our understanding and interpretation of gene expression patterns in cells and tissues. We argue that defining gene expression patterns at the subcellular level is crucial to 1) understand the differences between the cytosolic and nuclear transcriptomes and how each fraction contributes to the global gene expression patterns, 2) investigate the dynamics of subcellular RNA localization between different cells and different developmental stages as a regulatory mechanism to control protein expression and influence RNA function, and 3) evaluate the reliability of using nuclear transcriptome in single-cell studies as a proxy for whole-cell analysis. In this study, we therefore aimed to obtain a comprehensive overview of the subcellular localization of the different RNA transcripts in the human brain.

One group of transcripts strongly associated with the cytosol was the nuclear-encoded-mitochondrial transcripts. The biogenesis of the mitochondria relies on dual genetic origins; the nuclear and the mitochondrial genomes, but 99% of the mitochondrial proteins are encoded in the nucleus ^34,40^. This imposes the need for tight coordination between the two compartments to regulate gene expression. The detailed mechanisms that regulate gene and protein expression coordination between the nucleus and the mitochondria are not well understood. Nevertheless, it is believed that this coordination occurs both at mRNA and protein levels. Previous reports demonstrated that many mRNAs encoding NEMPs are localized to the outer membrane of mitochondria and it is believed that these mRNAs are translated locally ^41^. It has also been demonstrated that mRNAs coding for NEMPs localize to and are translated by ribosomes in the cytoplasm ^42^. However, an accurate estimation of the fraction of NEMP mRNAs that localize to the mitochondrial-membrane and cytosol is still an open question, particularly in humans as most of the current knowledge thus far is based on drosophila and yeast ^43^. In this study, we demonstrate, for the first time, that NEMP mRNAs in the human brain are significantly enriched in the cytosol as compared to the rest of the protein-coding genes. We validated this observation in ENCODE cell lines, where independent extraction protocols were employed. We also showed that the cytosolic localization of NEMP mRNAs was not a result of their association to the mitochondrial membrane since the mitochondria end up in the nuclear fraction in our cellular fractionation procedure. NEMPs have previously been assigned to classes based on their translation location in relation to mitochondria (MLR class) ^43^, but we found no correlation between increased cytosolic localization and MLR class (data not shown). We were also unable to find any correlation with the strength of the mitochondrial localization signal or scores based on evidence for mitochondrial localization (TargetP score or MitoCarta2.0 score) from the MitoCarta database ^34^ (data not shown). We further determine that NEMP mRNAs on average have a longer half-life than the rest of protein-coding genes. Thus, our data suggest that NEMP mRNAs are rapidly transported from the nucleus after transcription and accumulate in the cytoplasm until their translation is required by the mitochondria. In line with our results, a recent study in yeast sought to monitor nuclear and mitochondrial gene expression during mitochondrial biogenesis. The study demonstrated that the synchronization between the two compartments does not occur on the transcriptional level. Instead, they are rapidly synchronized on protein translation level ^44^

It is becoming increasingly evident that subcellular localization of lncRNAs is strongly associated with their function. In light of this, ongoing efforts such as the LncATLAS and RNALocate are aiming to establish a complete map for subcellular localization of lncRNAs in various tissue types ^3,29^. So far, these databases rely on data from cell lines and lack knowledge about the subcellular localization of lncRNA in tissues and during development. Studies that have investigated cellular localization of lncRNAs during development were mainly performed in drosophila and zebrafish ^45,46^. Therefore, in this study, we sought to provide global data of lncRNA subcellular localization in the human brain for two developmental stages. In accordance with previous studies, we found that the majority of lncRNAs are more abundant in the nucleus than in the cytosol ^22^. We further validated previous findings of tissuespecific expression and subcellular localization of lncRNAs in the adult frontal cortex, fetal frontal cotex, and cerebellum. The higher number of cytosolic and nuclear-enriched lncRNAs detected in the adult frontal lobe compared to the fetal samples is most probably due to the larger number of adult brain samples used in this study, as well as a more homogeneous expression in the adult brain compared to the fetal samples (represented by 14 and 38-week embryos), providing better power to detect significant differences in localization

Considering the differences in subcellular localization between the different brain samples, we investigated whether subcellular localization of lncRNAs could vary between different tissue contexts. When we compared the subcellular localization patterns of lncRNAs in the brain samples and ten cell lines from the ENCODE project, we clearly show that some lncRNAs have opposite subcellular localization patterns in different samples. This indicates that the subcellular localization of certain lncRNAs is dynamic, which could be explained by several processes including tissue-specific functionality or regulation, or that some lncRNA are retained in the nuclear compartment to regulate their function, assuming that their site of action is in the cytosol. Our results provide a resource for lncRNAs enriched in cytosol and nucleus in both fetal and adult human brain samples. We also provide evidence for the differential subcellular localization of lncRNAs in tissues. These data further encourage the need to understand the mechanisms that control the subcellular fate of lncRNAs and the biological consequences of this process.

The impact of using transcriptome data obtained from either cytosolic or nuclear RNA as a reliable proxy for gene expression analysis is an active topic of investigation. Recently, single nucleus RNA sequencing has been introduced as an alternative to whole single-cell sequencing, particularly in tissues where it is challenging or impossible to recover intact cells ^36,47,48^. In addition to the data presented in this study, several other studies demonstrated significant differences between the nuclear and the cytosolic transcriptome ^17,49^. Also, the compartmentalization of mRNA transcripts in the nucleus was shown to play an active role in regulating gene expression in different biological contexts and to prevent fluctuations arising from bursts in gene transcription ^12,50^. On the same note, several studies revealed a higher correlation between RNA-seq data from the cytosol and whole-cell RNA-seq compared with RNA-seq data from nuclei ^15,50^. On the other hand, several studies comparing data from single-nuclei and single-cell concluded that nuclear RNA sequencing faithfully replicates data from the whole-cell ^36,39,47^. In light of this, we sought to analyze gene expression differences between fetal and adult brain cortex, and compare the results from cytosol vs cytosol RNA-seq to the results from nucleus vs nucleus RNA-seq. We identified almost twice the number of differentially expressed genes in the cyto vs cyto analysis compared to the nuc vs nuc analysis. Our data show that even though there was an overlap between the lists of DEGs between two analyses, almost 50% of the cyto vs cyto and 35% of the nuc vs nuc DEGs were unique for each analysis. We also note that 61 transcripts showed contradictory differential expression patterns between the two analyses.

A recent study by Lake *et al*. ^39^ found high concordance between single nuclei and whole-cell data and concluded the efficiency of single nuclei sequencing to interpret cellular diversity. We found on average more than 50% of cytosolic-enriched and nuclear-enriched transcripts from their study overlap with cytosolic and nuclear-enriched transcripts found in our data. We also found overlaps, particularly for the metabolic processes, between the two data sets for the list of ontology categories enriched in both the cytosol (or whole cell in the Lake et al. study) and nucleus. While we find a large overlap in genes identified as differentially expressed between samples using comparisons of cytosolic vs cytosolic and nuclear vs nuclear RNA, respectively, our data also highlight that certain categories of genes will give rise to differences in results depending on the cellular compartment used. This includes lncRNA, which are generally overrepresented in the nucleus, but primarily thought to be located in the compartment where they are functional. We further find many genes to be exclusively found in either cyto vs cyto or nuc vs nuc analysis, with e.g. differential expression of genes involved in translational processes and ribosomal function to be better assayed using cytosolic RNA. In single-cell analysis, there is now a range of protocols ranging from those employing very mild lysis that will primarily interrogate cytosolic RNA, to harsh lysis and single nucleus sequencing that will capture the whole cell or only nuclear RNA, respectively. Our results show important differences that may arise depending on the protocol used, and highlight the importance of which subcellular fraction that is analyzed in interpreting and comparing results from single-cell studies.

## Conclusions

Our data provide the first resource for the subcellular localization of RNA transcript in the human brain. We show significant differences in RNA expression for both protein-coding and lncRNAs between the cytosol and the nucleus. We also provide novel knowledge about the subcellular localization of NEMP transcripts, which could provide deeper insights into mitochondrial gene expression regulation. Our data suggest that using cytosol or nuclear RNA as a source for gene expression analysis might bias measurements of transcript levels.

## Methods

### Sample preparation

Cytoplasmic and nuclear RNA was purified from brain samples and SHSY-5Y cells using Cytoplasmic and nuclear RNA purification kit (Norgen) with modifications as published in ^28^. In short, Cytoplasmic and nuclear RNA was purified from two fetal frontal cortex using Cytoplasmic and nuclear RNA purification kit (Norgen) with modifications as illustrated in Figure 1. In short, 20 mg of frozen tissues were ground in liquid nitrogen using mortar and pestle. Tissue powder was transferred to ice cold 1.5 ml tubes. Then, 200 μl lysis buffer (Norgen) was added to the grounded tissue. The tubes were incubated on ice for 10 minutes and then centrifuged for 3 minutes at 13,000 RPM to separate the cellular fractions.

The supernatant containing the cytoplasmic fraction and the pellet containing the nuclei were mixed with 400 μl 1.6 M sucrose solution and carefully layered on the top of two 500 μl sucrose solution in two separate tubes. Both fractions were then centrifuged on 13,000-RPM for 15 minutes (4C°). The cytoplasmic fraction was collected from the top of the sucrose cushion and the cytoplasmic RNA was then further purified according to Norgen kit recommendations. The nuclear pellet was collected from the bottom of the tube and washed with 200 μl 1× PBS. The nuclei were collected after another centrifugation at 13,000 RPM for 3 minutes. The nuclear RNA was purified from the nuclear fraction according to the Norgen kit recommendations.

Prior to RNA extraction, SHSY-5Y cells were cultured in 1:1 mixture of Ham’s F12 and DMEM (Gibco) medium (Gibco) with L-glutamine and phenol red, supplemented with 2 mM l-glutamine (Sigma), 10% FBS (Sigma) and 1× PEST (Sigma). Cells were incubated at 37 °C, 5% CO2. 3*10^7^ cells were used for RNA purification.

### RNA sequencing

Only cytoplasmic RNA was treated with the Ribo-Zero Magnetic Gold Kit for human/mouse/Rat (Epicentre) to remove ribosomal RNA, and purified using Agencourt RNAClean XP Kit (Beckman Coulter). The rRNA depleted cytoplasmic RNA and nuclear RNA were then treated with RNaseIII according to Ion Total RNA-Seq protocol v2 and purified with Magnetic Bead Cleanup Module (Thermo Fisher). Sequencing libraries were prepared using the Ion Total RNA-Seq Kit for the AB Library Builder™ System and quantified using the Fragment analyzer (Advanced Analytical). The quantified libraries were pooled followed by emulsion PCR on the Ion OneTouch 2 system and sequenced on the Ion Proton System.

### Read mapping and expression quantification

Data was delivered as BAM files from TMAP, but since these were not aligned with an aligner for spliced reads, therefore, the BAM files were converted into FASTQ format. Quality metrics of the raw data were determined with RSeqC 2.6.1 ^51^. Reads shorter than 36bp were filtered out and adaptor sequences were trimmed from the remaining data using cutadapt v1.9.1. Read alignment was done with STAR v2.4.2 ^52^, with default parameters using the human reference genome (hg19). Gene expression was quantified from the alignments using HTSeq count (v0.6.1) ^53^ by providing Ensembl gene annotation (v75) in GTF format. Only uniquely mapped reads overlapping with exonic regions of a single gene were considered for quantification. Gene biotype annotation was extracted from the Ensembl GTF file.

### Differential expression analysis

Strand-specific read counts were provided to R package DESeq2 (v1.12.4) ^54^ to identify differentially expressed genes (adjusted *p-value* <0.05, Benjamini-Hochberg correction). Gene counts were normalized prior to differential analysis using the normalization method implemented in DESeq2. To identify differentially expressed snoRNA genes, all the rRNA and tRNA genes were removed from the quantified genes prior to performing differential analysis. For the cyto-cyto and nuc-nuc differential gene expression analysis, we subsampled the read counts from the samples to obtain equal library sizes using metaseqR ^55^. Differential expression analysis was performed in the similar way as mentioned earlier.

### Samples clustering and PCA analysis

Euclidean distances between the samples were calculated on regularized log-transformed (rlog) counts. The distance matrix was used to compute hierarchical clustering of the samples. The Principal Component Analysis (PCA) was performed with prcomp function in R using rlog values of top 500 genes with high variance. Clustering and PCA were based on protein-coding and lncRNA genes.

### Gene Ontology enrichment

The gene ontology (GO) term enrichment of DEGs was analyzed using GOSeq (v.1.24) ^56^, controlling for gene-length bias. All genes were used as a background set in order to find over-represented GO terms in the DEGs. Hypergeometric test (Benjamini and Hochberg correction) was performed to compensate for multiple testing at a significance level set to a value of <0.05 in our analyses.

### Gene length distribution

Genes expressed in all fetal and adult tissues (base mean expression >= 10) were sorted by length. We then employed a sliding window comprised of 200 genes in steps of 40 genes. The log2-fold-change values for the 200 genes within each length bin were averaged and standard errors (SE) for a bin were calculated by propagating the SE determined from the bin’s log2-fold-change values and the mean SE of the individual genes reflecting their sample variability. Similarly, we computed the relationship of transcript and UTRs length, and gene length with expression (RPKM).

### Nuclear-encoded-mitochondrial proteins

A list of 1,158 human Nuclear-encoded-mitochondrial proteins (NEMPs) was obtained from the Mitocarta2.0 project (https://www.broadinstitute.org/pubs/MitoCarta/). Expression fold changes were estimated after normalizing the samples for only protein-coding genes using DESeq2. Significance of difference in fold change distribution between NEMPs and other protein-coding genes (baseMean >= 10) was determined using the nonparametric Mann-Whitney-Wilcoxon test.

### ENCODE data

Gene counts (hg19) of polyA and non-polyA RNAseq data of cytoplasmic and nuclear fractions for the cell lines: GM12878, HUVEC, HepG2, HeLa-S3, NHEK, K562, IMR90, MCF-7, A549, and SK-N-SH, were obtained from ENCODE project (GEO:GSE3056). For NEMPs and ribosomal protein-coding genes analyses, polyA data were normalized for protein-coding genes. For lncRNA analysis, gene counts (polyA) were normalized for protein-coding and lncRNA genes.

### RNA FISH

RNA FISH assays were performed using a custom-designed oligonucleotide probe set for OAT and SLC25A5, both labeled with CAL Fluor^®^ Red 610 and DesignReady Probe Sets for MALAt1 labeled with Quasar^®^ 670 Dye, and GAPDH labeled with CAL Fluor^®^ Red 610 (Stellaris^®^, LGC Biosearch Technologies). 48 oligonucleotides were selected for OAT, MALAT1, and GAPDH, and 27 oligonucleotides for SLC25A5. All probes were designed with a minimal mismatch.RNA FISH was carried using hybridization and wash buffers from LGC Biosearch Technologies. In short, 10^5 human SHSY-5Y cells seeded on 18 mm #1 round cover glass (VWR) in a 12 well culture plate. Cells were cultured in 1:1 mixture of Ham’s F12 and DMEM (Gibco) medium (Gibco) with L-glutamine and phenol red, supplemented with 2 mM l-glutamine (Sigma), 10% FBS (Sigma) and 1× PEST (Sigma). Cells were maintained in culture at 37□°C and 5% CO2. Following overnight incubation, cells were fixed and permeabilized according to manufacturer recommendations. Cells were incubated for 16 hours with FISH probes in the hybridization buffer (LGC Biosearch Technologies). Cells were then washed using wash buffer A (LGC Biosearch Technologies) for 30 min at 37□°C, and then DNA staining with DAPI (1□μg/ml) for 30 min at 37□°C. Finally, cells were washed in wash buffer B, and coverslips were mounted in Vectashield anti-fade (VectorLabs) and imaged immediately. Imaging was carried out on Zeiss Axio Imager Z2 Widefield Fluorescence Microscopy using an Olympus 63x/1.42 PlanApo objective and using and ZEN imaging software. Z-stacks were collected at 0.5 □ μm intervals for 18 fields per sample. Z-stacks were deconvolved using orthogonal projection (ZEN imaging software)

## Declarations

### Ethics approval and consent

All samples were collected with informed consent and the study was approved by the regional ethical review board in Uppsala (dnr 2012/082).

### Availability of data and material

RNA-seq data are deposited in the NCBI Sequence Read Archive (SRA) under accession GSE110727).

### Competing interests

The authors in this study declare no competing interest

### Authors’ contributions

A.Z. and L.F. conceived and designed the study. A.Z., A.N., Å.B., J.W, A.A and L.F. coordinated experiments and analysis. A.N., Å.B conducted the bioinformatics analysis. A.Z. did the sample preparation and experimental analysis. All authors participated in discussions of different parts of the study. A.Z., A.N., Å.B., and L.F. wrote the manuscript. All authors read and approved the manuscript.

## Acknowledgments

The authors would like to acknowledge support from Science for Life Laboratory, the National Genomics Infrastructure, NGI, and Uppmax for providing assistance in massive parallel sequencing and computational infrastructure. We are also grateful for the Bioinformatics Long-term Support from NBIS (SciLifeLab) for this project. This work was supported by grants Swedish Research Council (2012-4530 and 2012-2639) and the European Research Council ERC Starting Grant Agreement n. 282330.

